# DNA methylation profiling of breast cancer cell lines along the epithelial mesenchymal spectrum - implications for the choice of circulating tumour DNA methylation markers

**DOI:** 10.1101/182600

**Authors:** Anh Viet-Phuong Le, Marcin Szaumkessel, Tuan Zea Tan, Jean-Paul Thiery, Erik W Thompson, Alexander Dobrovic

## Abstract

Epithelial-mesenchymal plasticity (EMP) is a dynamic process whereby epithelial carcinoma cells reversibly acquire morphological and invasive characteristics typical of mesenchymal cells, which facilitates metastasis. Understanding the methylation differences between epithelial and mesenchymal states may assist in the identification of optimal DNA methylation biomarkers for the blood-based monitoring of cancer. Methylation-sensitive high-resolution melting (MS-HRM) was used to examine the promoter methylation status of a panel of established and novel markers in a range of breast cancer cell lines spanning the epithelial-mesenchymal spectrum. Pyrosequencing was used to validate the MS-HRM results. The results indicate an overall distinction in methylation between epithelial and mesenchymal phenotypes. The mesenchymal expression markers *VIM, DKK3* and *CRABP1* were methylated in the majority of epithelial breast cancer cell lines while methylation of the epithelial expression markers *GRHL2, MIR200C* and *CDH1* was restricted to mesenchymal cell lines. We also examined EMP association of several methylation markers that have been used to assess minimal residual disease. Markers such as *AKR1B1* and *APC* methylation proved to be selective for epithelial breast cell lines, however *RASSF1A, RARß, TWIST1* and *SFRP2* methylation was seen in both epithelial and mesenchymal cell lines, supporting their suitability for a multi-marker panel.

## Background

Analysis of circulating tumour DNA (ctDNA) is increasingly being used for monitoring of minimal residual disease (MRD) during cancer treatment. Detection of ctDNA is dependent on the use of cancer-specific markers [1]. As breast cancer has few recurrent single site mutations outside of those in the *PIK3CA* gene [2], attention is being shifted to DNA methylation in which a relatively small panel of markers might be applicable to the majority of tumours. As DNA methylation markers have been reported to differ across the epithelial-mesenchymal spectrum, identification of a panel of DNA methylation markers that take epithelial-mesenchymal plasticity (EMP) into account would be desirable. Several groups have performed DNA methylome-wide analysis to compare the methylation profiles in relation to EMT status, in lung cancer cell lines [3], ovarian cancer cell lines [4], and breast cancer cell lines [5, 6].

Epithelial-mesenchymal plasticity (EMP) refers to dynamic cellular transformation across the epithelial-mesenchymal axis. This is a crucial event during normal development, and one of the mechanisms that cancer cells take advantage of in order to disseminate [7]. Epithelial carcinoma cells acquire genetic and epigenetic alterations, changing their morphology and behaviour to become more migratory and invasive. Through epithelial-mesenchymal transition (EMT), these cells can escape from the tumour into the tissues and circulation, and travel to secondary sites, where they undergo mesenchymal-epithelial transition (MET) to allow formation of metastases [8]. Tumours and their metastases have been shown to consist of heterogeneous mixtures of epithelial and mesenchymal cells [9, 10].

The understanding of DNA methylation in EMP, especially in breast cancer, remains elusive. Thus, we sought to examine how the methylation of both established and novel markers related to the epithelial and mesenchymal states across a panel of breast cancer cell lines that were positioned across the epithelial–mesenchymal spectrum. Our hypothesis was that relating the locus-specific methylation status with epithelial and mesenchymal states would enable clarity in the choice of DNA methylation markers for monitoring minimal residual disease (MRD) using ctDNA. To our knowledge, this is the first study that characterises DNA methylation status of a large panel of breast cancer cell lines spanning the epithelial–mesenchymal spectrum. Examining DNA methylation across the epithelial-mesenchymal spectrum is a novel approach to identify optimal DNA methylation markers for ctDNA.

## Materials and methods

### Breast cancer cell lines

Frozen cell pellets of the following breast cancer cell lines, MDA-MB-231, BT-549, HCC1569, HCC1806, SK-BR-3, HCC70, MDA-MB-468, MDA-MB-453, MCF-7, HCC1954, HCC1419, T-47D, BT-20, BT-474, CAL-120, CAL-148, MCF 10A, SUM-149PT, SUM-159PT, UACC-732, UACC-893, ZR-75-1, MFM-223, and Hs578T were used to extract DNA for methylation analysis. These cell pellets were kindly provided by Riley Morrow and Tracy Cardwell from the Olivia-Newton John Cancer Research Institute, from cells that were originally from ATCC and/or had been STR authenticated. DNA from PMC42-ET and PMC42-LA cells was kindly provided by Dr. Mark Waltham, St. Vincent’s Hospital, Melbourne.

### DNA extraction from cell pellets

DNA was extracted from the pellets using the DNeasy^®^ Blood & Tissue Kit (Qiagen, Hilden, Germany) according to the manufacturer’s instructions. Briefly, in a screw cap tube (Neptune), 200 μl of 1 x PBS and subsequently 20 μl of Proteinase K (Scimar, Cat no. LS004224) were added to each cell pellet, which contained less than 5 x 10^6^ cells. The mixtures were vortexed briefly for 10 seconds. 200 μl of Buffer AL was then added into each cell pellet mixture. The mixtures were pulse vortexed for 15 seconds and briefly centrifuged, followed by incubation in an oven at 56˚C overnight. The tubes containing cells and buffers were taken out of the oven and cooled for 5 minutes at room temperature before DNA extraction and clean up. The tubes were vortexed and briefly centrifuged. 200 μl of ethanol (Sigma-Aldrich) was subsequently added into each tube and the mixtures were pulse vortexed and centrifuged. The contents in the screw cap tubes were carefully transferred into spin columns provided in the Blood & Tissue Kit, which were then centrifuged at 8,000 rpm for 1 minute. The flow through and collection tubes were discarded and new collection tubes were used for the subsequent washing steps. The spin columns were washed by adding 500 μl of Buffer AW1 and centrifuging at 8,000 rpm for 1 minute. Next, 500 μl of Buffer AW2 was added and the columns were centrifuged at 8,000 rpm for 1 minute. The spin columns were then centrifuged at 14,000 rpm for 3 minutes without adding any more washing buffer. During washing steps using the spin columns, the flow through and collection tubes were discarded and the spin columns were inserted into new collection tubes for each step. To elute DNA, 50 μl of Buffer AE, which was pre-warmed at 72˚C, was added directly into each spin column. After 5 minutes of incubation at room temperature, the columns were centrifuged at 14,000 rpm for 3 minutes. The DNA was then quantified using the DS-11 Spectrophotometer (DeNovix, Wilmington, DE) and stored at 4˚C for short-term storage and at -20˚C for long-term storage.

### Bisulfite conversion

Bisulfite conversion was performed with the EZ DNA Methylation-Lightning kit (Zymo Research, Irvine, CA). 500-1,000 ng of DNA was modified using the bisulfite kit according to the manufacturer’s instructions. In 8-tube strips, PCR-grade water was added to the required volume of DNA to make up a total volume of 20 μl for each bisulfite converted reaction. Next, 130 μl of Lightning Conversion Reagent was added to each of the DNA solution, followed by pipetting up and down five times. After being centrifuged, the 8-tube strips were placed into a C1000 Touch^™^ thermal cycler (Bio-Rad, USA). The step-by-step cycling program was as followed: (1) denaturation step at 98˚C for 8 minutes, (2) incubation step at 54˚C for 60 minutes, (3) hold at 4˚C for up to 20 hours.

After the conversion, the bisulfite converted DNA was cleaned up by transferring the DNA in the 8-tube strips to the Zymo-Spin™ IC columns containing 600 μl of M-binding buffer. The columns were inverted 5 times to mix the contents, and subsequently centrifuged at 14,000 rpm for 1 minute. The flowthrough was discarded and the spin columns were placed back into the collection tubes. 100 μl of M-washing buffer was added to each spin column, which was centrifuged at 14,000 rpm for 1 minute. Next, 200 μl of L-Desulfonation buffer was added to each spin column, followed by a 20-minute incubation at room temperature. The spin columns were centrifuged at 14,000 rpm for 1 minute. The columns were washed by the addition of 200 μl of M-Wash buffer and centrifugation at 14,000 rpm for 1 minute. This step was repeated once. After two washing steps, the spin columns were placed into new Eppendorf tubes. After adding 10 μl of M-Elution buffer directly into each spin column, the columns were incubated for 5 minutes at room temperature and then centrifuged at 14,000 rpm for 1 minute. Another 10 μl of M-Elution buffer was added onto the spin columns, followed by 5-minute incubation at room temperature and 14,000 rpm centrifugation for 1 minute. A total of 20 μl of bisulfite converted DNA was obtained. Bisulfite converted DNA was stored at 4˚C for short-term storage and at -20˚C for long-term storage (longer than two months).

### Methylation-sensitive high resolution melting

MS-HRM was performed on a Rotor-gene 6000 (Corbett, Sydney, Australia) or a CFX Connect Realtime System (BioRad, Hercules, USA). For each MS-HRM run, a methylation standard series including 100%, 50%, 10%, 3% methylated and fully unmethylated standards was included as a guide for detecting methylation and for calling methylation levels [11, 12]. Each sample was run in duplicate. MS-HRM primers were designed following the guidelines described previously [13, 14]. Reverse MS-HRM primers were biotinylated to allow the subsequent bisulfite pyrosequencing. The 20 μl of PCR reaction mix consisted of 1 x PCR buffer (Qiagen), 1.5 to 3.0 mM MgCl2, 200uM dNTP mix (Fisher Biotech, Perth, Australia), 200-400 nM forward and reverse primers, 1 x SYTO9 intercalating dye (Thermo Fisher Scientific, Waltham, MA), 0.5U HotstarTaq polymerase (Qiagen) and 10 ng of bisulfite converted DNA (theoretical amount from the assumption of no DNA loss during bisulfite conversion). The primer sequences and the PCR conditions for all MS-HRM assays are listed in Table S1 in Additional file 1 and Table S2 in Additional file 2, respectively.

The levels of homogeneous methylation were stated numerically as percentage of methylation. Whereas the estimated levels of heterogeneous methylation were described as ‘very high’, ‘high’, ‘moderate’, ‘low’ and ‘very low’ for very high, high, moderate, low and very low levels of methylation, respectively, based on the extent that the melting profiles extend to the methylated area.

### Bisulfite Pyrosequencing

Bisulfite pyrosequencing was used subsequently to MS-HRM in order to quantify the methylation percentage at each CpG position [12]. The MS-HRM primers were used for bisulfite pyrosequencing. The reverse primers were biotinylated and forward primers were used as sequencing primers. Representative samples from *RASSF1A, RARβ, AKR1B1, GRHL2, CRABP1, SFRP2* and *GFRA1* previous assessment by MS-HRM were pyrosequenced. Methylation standards were also included in pyrosequencing reactions. Two μl of PCR products from MS-HRM assays were used as templates for pyrosequencing.

The pyrosequencing reaction was performed on a Qseq pyrosequencer (Bio Molecular Systems, Sydney, Australia) using the Pyro Gold Q24 reagents (Qiagen). The injector set-up, priming and testing were performed according to the Qseq instructions. After inserting the 48-well disc into the Qseq pyrosequencer, 8 μl of MilliQ water, 1 μl of well mixed Streptavidin Mag Sepharose beads and 2 μl of PCR templates was loaded into each well. Denaturation of templates and subsequent annealing of biotinylated single stranded DNA into beads were performed after the run was started. 2 μl of the pyrosequencing primers (5uM) were used for each reaction. Table S3 in Additional file 3 provides the sequences before bisulfite treatment, sequences for analysis and dispensation order for each assay. Qseq advanced software (Bio Molecular Systems, v2.0.11) was utilized to set up CpG assays (CpG mode) and run files. The obtained data were analysed and quantified with the Qseq Advanced Software.

## Results

### Ranking of breast cancer cell lines across the epithelial-mesenchymal spectrum

In order to relate the DNA methylation status to epithelial and/or mesenchymal phenotypes, we first ranked the twenty-six breast cancer cell lines on the epithelial-mesenchymal spectrum using a previously published quantitative scoring system [15]. Epithelial cell lines have negative scores on this scale, with MFM-223 being the most epithelial of the cell lines (Figure 1). Positive scores indicate mesenchymal cell lines, with Hs578T being the most mesenchymal of the cell lines (Figure 1). The MDA-MB-468, HCC1954, HCC70, BT-20, HCC1806, HCC1569, SUM-149PT, and CAL-120 cell lines with intermediate scores were considered to have intermediate epithelial-mesenchymal phenotypes [15].

**Figure 1:**
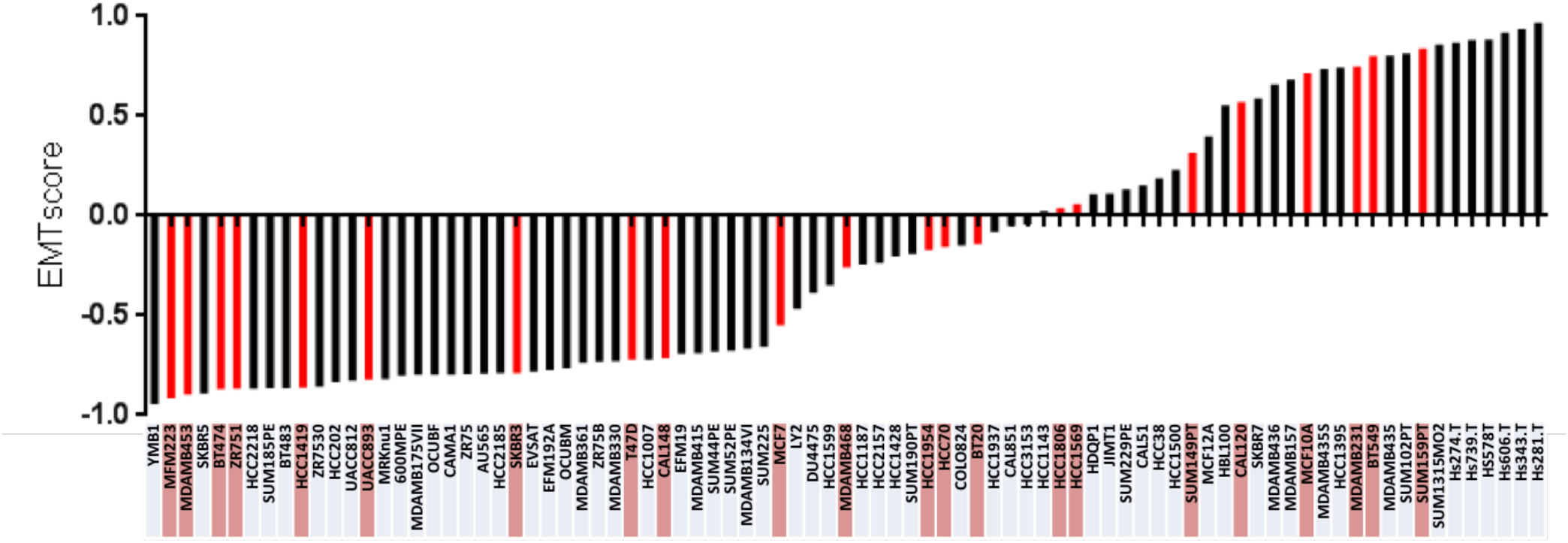
Breast cancer cell lines with available EMT score are highlighted and placed from the most epithelial to the most mesenchymal in the epithelial mesenchymal spectrum.

Data to rank the PMC42 cell line (PMC42-ET) and its more mesenchymal sub-line (PMC42-LA) [16] were not available. PMC42-LA and PMC42-ET were both placed within the Basal B group based on independent clustering of gene expression data (Blick, Tomaskovic-Crook, Neve, Thompson; unpublished data).

Table 1 summarises the hormonal receptor status, vimentin status, molecular subgroup, and epithelial and mesenchymal status of these cell lines based on publically available datasets and publications. The data for expression of estrogen receptor (ER), progesterone receptor (PR) and human epidermal growth factor receptor 2 (HER2) of selected cell lines were found in the American Type Culture Collection (ATCC) website and references therein [17]. For the cell lines whose hormone receptor status could not be found in the ATCC, this information was extracted from Kao *et al*. [18] and Lehmann *et al*. [19], as indicated in Table 1. Vimentin expression was previously determined by us [20, 21]. Breast cancer cell line intrinsic subgroups, including luminal, basal A and basal B, were according to the Neve *et al*. [22], Hoeflich *et al*. [23] and Heiser *et al*. [24] gene expression studies. HER2-amplification status in MDA-MB-453, BT-474, HCC1419, UACC-893, HCC1954, HCC1569 and SK-BR-3 cell lines, reported in the GSE12790 data by Hoeflich *et al*. [23], was confirmed by Kalous *et al*. using fluorescence *in situ* hybridization (FISH) [25]. We observed that the majority of luminal and HER2-amplified cell lines are epithelial, whereas basal B and triple negative (ER-/PR-/HER2-) cell lines are mesenchymal. Basal A cell lines usually have an intermediate phenotype, regardless of HER2 status.

**Table 1:**
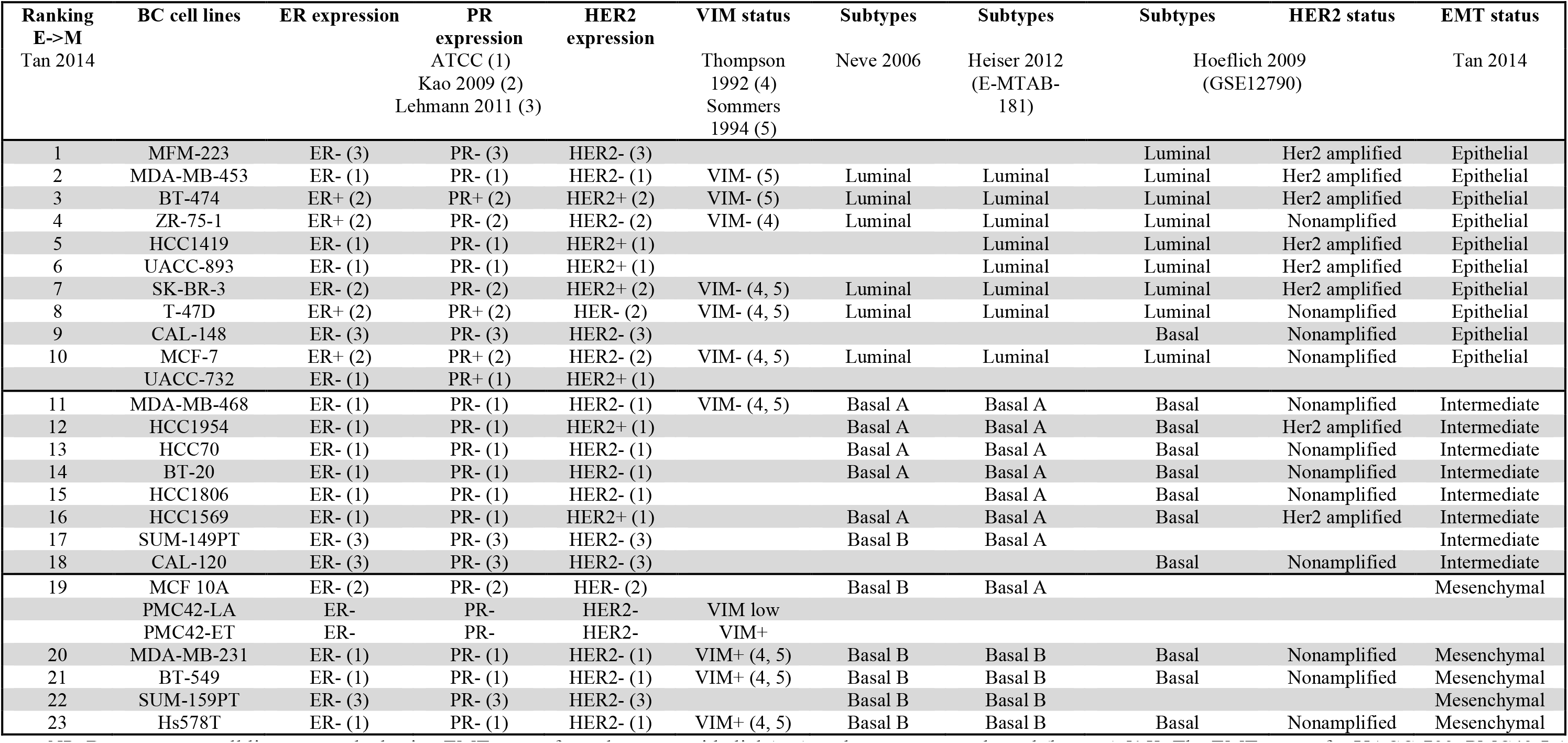
Summary of hormonal receptor status, VIM status, molecular subtypes, and EMT status of a panel of 26 breast cancer cell lines according to published literature.

| Ranking<br>E->M<br>Tan 2014 | BC cell lines | ER expression | PR<br>expression<br>ATCC (1)<br>Kao 2009 (2)<br>Lehmann 2011 (3) | HER2<br>expression | VIM status<br>Thompson<br>1992 (4)<br>Sommers<br>1994 (5) | Subtypes<br>Neve 2006 | Subtypes<br>Heiser 2012<br>(E-MTAB-181) | Subtypes<br>Hoeflich 2009<br>(GSE12790) | HER2 status | EMT status<br>Tan 2014 |
| --- | --- | --- | --- | --- | --- | --- | --- | --- | --- | --- |
| 1 | MFM-223 | ER- (3) | PR- (3) | HER2- (3) |  |  |  | Luminal | Her2 amplified | Epithelial |
| 2 | MDA-MB-453 | ER- (1) | PR- (1) | HER2- (1) | VIM- (5) | Luminal | Luminal | Luminal | Her2 amplified | Epithelial |
| 3 | BT-474 | ER+ (2) | PR+ (2) | HER2+ (2) | VIM- (5) | Luminal | Luminal | Luminal | Her2 amplified | Epithelial |
| 4 | ZR-75-1 | ER+ (2) | PR- (2) | HER2- (2) | VIM- (4) | Luminal | Luminal | Luminal | Nonamplified | Epithelial |
| 5 | HCC1419 | ER- (1) | PR- (1) | HER2+ (1) |  |  | Luminal | Luminal | Her2 amplified | Epithelial |
| 6 | UACC-893 | ER- (1) | PR- (1) | HER2+ (1) |  |  | Luminal | Luminal | Her2 amplified | Epithelial |
| 7 | SK-BR-3 | ER- (2) | PR- (2) | HER2+ (2) | VIM- (4, 5) | Luminal | Luminal | Luminal | Her2 amplified | Epithelial |
| 8 | T-47D | ER+ (2) | PR+ (2) | HER- (2) | VIM- (4, 5) | Luminal | Luminal | Luminal | Nonamplified | Epithelial |
| 9 | CAL-148 | ER- (3) | PR- (3) | HER2- (3) |  |  |  | Basal | Nonamplified | Epithelial |
| 10 | MCF-7 | ER+ (2) | PR+ (2) | HER2- (2) | VIM- (4, 5) | Luminal | Luminal | Luminal | Nonamplified | Epithelial |
|  | UACC-732 | ER- (1) | PR+ (1) | HER2+ (1) |  |  |  |  |  |  |
| 11 | MDA-MB-468 | ER- (1) | PR- (1) | HER2- (1) | VIM- (4, 5) | Basal A | Basal A | Basal | Nonamplified | Intermediate |
| 12 | HCC1954 | ER- (1) | PR- (1) | HER2+ (1) |  | Basal A | Basal A | Basal | Her2 amplified | Intermediate |
| 13 | HCC70 | ER- (1) | PR- (1) | HER2- (1) |  | Basal A | Basal A | Basal | Nonamplified | Intermediate |
| 14 | BT-20 | ER- (1) | PR- (1) | HER2- (1) |  | Basal A | Basal A | Basal | Nonamplified | Intermediate |
| 15 | HCC1806 | ER- (1) | PR- (1) | HER2- (1) |  |  | Basal A | Basal | Nonamplified | Intermediate |
| 16 | HCC1569 | ER- (1) | PR- (1) | HER2+ (1) |  | Basal A | Basal A | Basal | Her2 amplified | Intermediate |
| 17 | SUM-149PT | ER- (3) | PR- (3) | HER2- (3) |  | Basal B | Basal A |  |  | Intermediate |
| 18 | CAL-120 | ER- (3) | PR- (3) | HER2- (3) |  |  |  | Basal | Nonamplified | Intermediate |
| 19 | MCF 10A | ER- (2) | PR- (2) | HER- (2) |  | Basal B | Basal A |  |  | Mesenchymal |
|  | PMC42-LA | ER- | PR- | HER2- | VIM low |  |  |  |  |  |
|  | PMC42-ET | ER- | PR- | HER2- | VIM+ |  |  |  |  |  |
| 20 | MDA-MB-231 | ER- (1) | PR- (1) | HER2- (1) | VIM+ (4, 5) | Basal B | Basal B | Basal | Nonamplified | Mesenchymal |
| 21 | BT-549 | ER- (1) | PR- (1) | HER2- (1) | VIM+ (4, 5) | Basal B | Basal B | Basal | Nonamplified | Mesenchymal |
| 22 | SUM-159PT | ER- (3) | PR- (3) | HER2- (3) |  | Basal B | Basal B |  |  | Mesenchymal |
| 23 | Hs578T | ER- (1) | PR- (1) | HER2- (1) | VIM+ (4, 5) | Basal B | Basal B | Basal | Nonamplified | Mesenchymal |

*NB*: Breast cancer cell lines are ranked using EMT score, from the most epithelial (top) to the most mesenchymal (bottom) [15]. The EMT scores for UACC-732, PMC42-LA and PMC42-ET are not available. Hormonal receptor status, including ER, PR and HER2 expression, were determined from (1) the ATCC website and references therein [17], (2) Kao 2009 [18] and (3) Lehmann 2011 [19]. VIM status from immunofluorescent analysis was reported in (4) Thompson 1992 [21] and (5) Sommers 1994 [20]. The VIM status and the hormonal receptor status of PMC42-LA and PMC42-ET are cited from [16] and unpublished data. Molecular subtypes were obtained from the studies of Neve 2006 [22], Heiser 2012 [24] using E-MTAB-181 dataset and Hoeflich 2009 [23] using GSE12790 dataset. The cell lines were also assigned the epithelial and mesenchymal status based on the study of Tan 2014 [15]. Unavailable information is left empty.

### The DNA methylation status of breast cancer cell lines

To obtain information about the methylation pattern and estimate the methylation level for the sixteen genes, we employed methylation-sensitive high resolution melting (MS-HRM), which amplifies regions of interest regardless of methylation status and enables the differentiation of methylated, unmethylated and partially methylated templates after the polymerase chain reaction (PCR) by high resolution melting (HRM) analysis [11, 13]. A series of methylation standards was included in each MS-HRM run to estimate the methylation level. By analysing the melting profiles of the amplified products from MS-HRM assays, homogeneous or heterogeneous methylation can be identified (Figure 2A and 2B).

**Figure 2:**
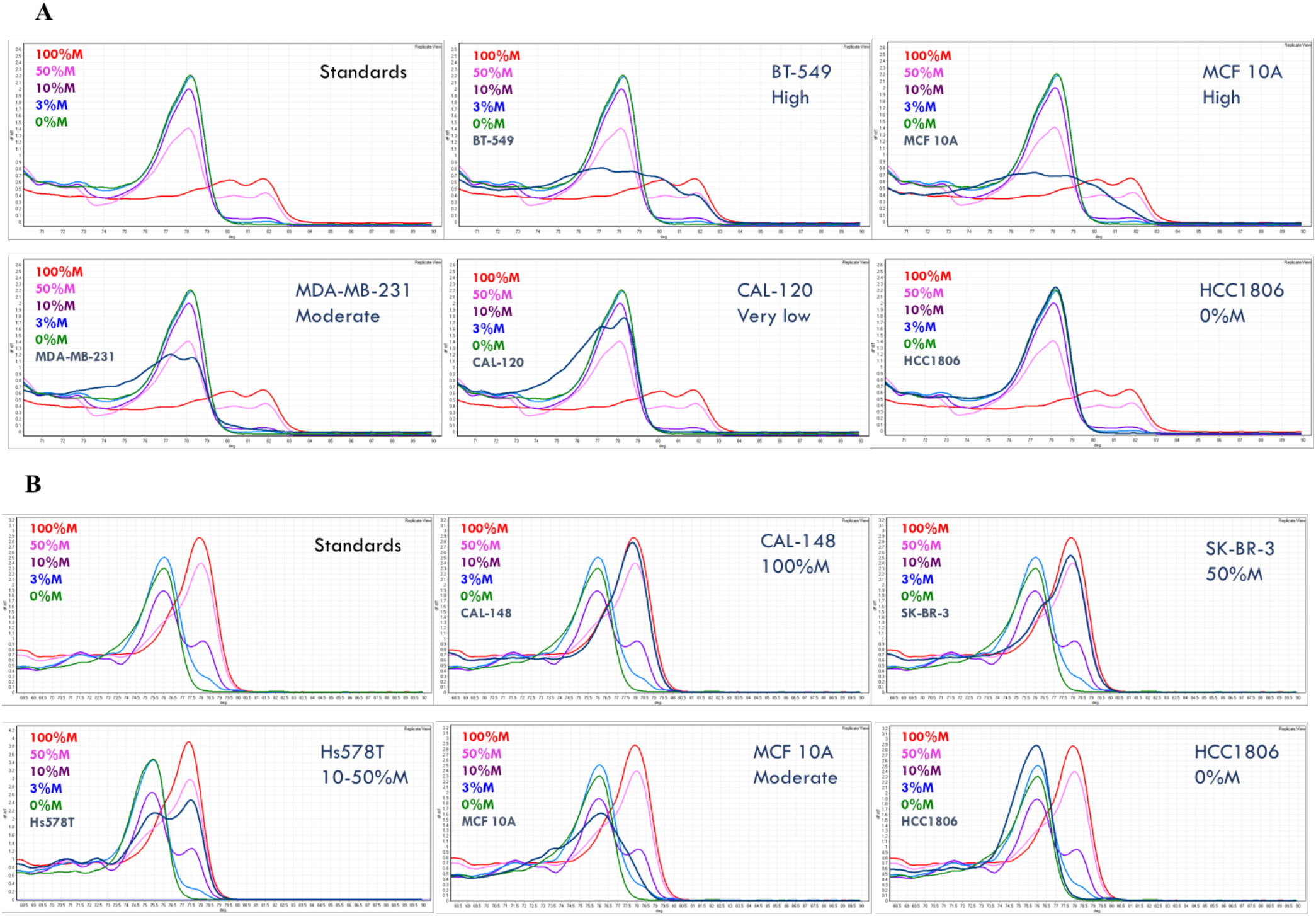
MS-HRM results of representative samples for (A) *GRHL2* and (B) *RASSF1A* show homogeneous methylation and heterogeneous methylation. Methylation level was numerical for samples with homogeneous methylation, whereas it was called ‘high, ‘moderate, ‘very low’ for high level, moderate level and very low level of heterogeneous methylation, respectively.

The melting profiles of the methylation standard series exemplify homogeneous methylation, in which CpG sites within the same amplicons are collectively methylated or unmethylated. The PCR products from the methylated templates with higher melting temperature have melt peaks to the right, while the PCR products from the unmethylated templates melt earlier, thus have melt peaks to the left. A sample with a melting profile resembling that of the methylation standards is homogeneously methylated. In this case, the methylation level can be semi-quantified using the methylation standard series. For example, HCC1806 is fully unmethylated for *GRHL2* (Figure 2A), CAL-148 is fully methylated and SK-BR-3 is homogeneously methylated at 50% for *RASSF1A* (Figure 2B).

When the methylation profile extends across either or both sides of the fully unmethylated control peak, the sample is heterogeneously methylated. Heterogeneous methylation patterns result as a consequence of heteroduplex formation from multiple different partially-methylated templates within the same sample [26]. Figure 2A gives examples of heterogeneous methylation in BT-549, MCF-10A, MDA-MB-231 and CAL-120 cell lines for *GRHL2*. Figure 2B shows an example of heterogeneous methylation for *RASSF1A* in the MCF 10A cell line.

In contrast to homogeneous methylation, the level of heterogeneous methylation is less readily determined by visual examination due to the complexity of the methylation patterns. We estimated the heterogeneous methylation level as very high, high, moderate, low or very low based on the degree to which the melting curves extend into the fully methylated profiles. The further the melting curves extend under the fully methylated peak, the higher the overall methylation levels (e.g. Figure 2A and 2B).

The PCR products were further assessed by bisulfite pyrosequencing to determine the average percentage at each CpG site for the selected markers, in order to obtain additional information about methylation, and to validate the MS-HRM results [12]. Due to the possible bias introduced by MS-HRM assays at given temperatures, (i.e., over-estimation or under-estimation of methylation levels), the methylation standards were also pyrosequenced together with the samples. Table 2 shows pyrosequencing data for the methylation standard series and representative samples of each methylation level calling for *GRHL2* methylation and *RASSF1A* methylation.

The pyrosequencing results for *GRHL2* (Table 2A) and *RASSF1A* (Table 2B) indicate that the estimation of methylation levels by MS-HRM is well matched with the values given by pyrosequencing. The degree of extension into the fully methylated profile is consistent with the levels of methylation overall. Pyrosequencing was also performed for other markers, including *AKR1B1, RARß, SFRP2, GFRA1* and *CRABP1* (Supplementary Figure S1 in Additional file 4). The concordance between the methylation level estimation and the pyrosequencing data of the samples relative to those of the methylation standard series supports our estimates of the methylation levels for heterogeneous methylation directly from the melting curves.

### Heterogeneous methylation is frequent for the studied markers

The methylation results for the studied markers in the breast cancer cell line panel are presented in Table 3. Samples with homogeneous methylation were marked with numerical values for their methylation levels, whereas those with heterogeneous methylation are marked as ‘very high’, ‘high’, ‘moderate’, ‘low’ and ‘very low’ as an estimation of their methylation levels, concordant with pyrosequencing results.

Some methylation markers commonly used for monitoring breast cancer, including *APC, RASSF1A* and *RARß*, predominantly displayed patterns consistent with homogeneous methylation. Other less commonly used markers for monitoring, such as *AKR1B1* and *SFRP2*, showed both homogeneous and heterogeneous methylation patterns in the cell line samples. In contrast, the majority of the other markers showed heterogeneous methylation patterns. This was most pronounced for *GRHL2, MIR200C* and *CDH1*. In addition, the majority of the mesenchymal cell lines displayed heterogeneous methylation patterns in most genes compared to the epithelial cell lines.

**Table 2:**
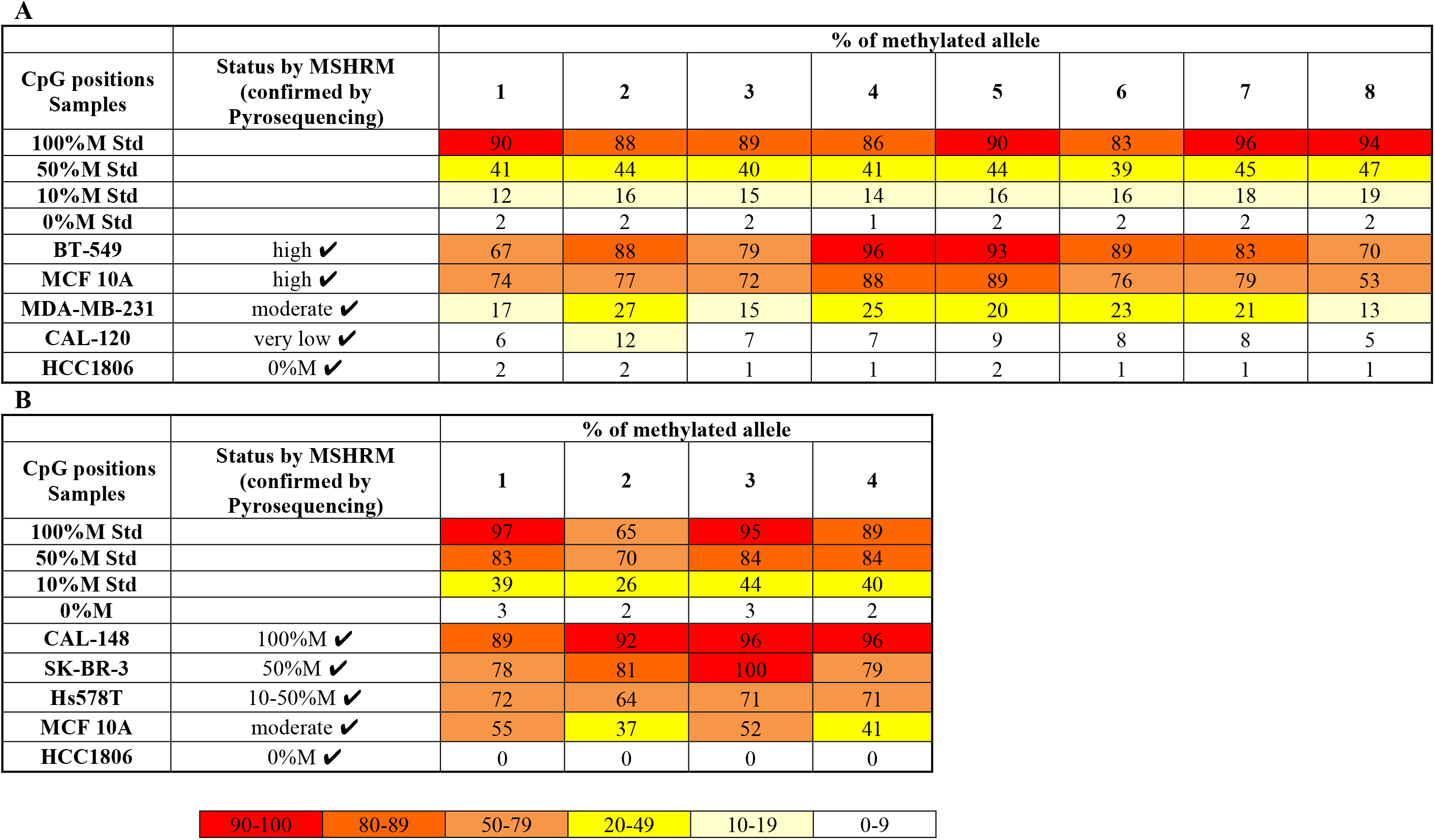
Pyrosequencing results of representative samples for (A) *GRHL2* and (B) *RASSF1A*. Pyrosequencing data validated the results from MS-HRM and provided complementary information on methylation level at each CpG position.

**Table 3:** Methylation data of the studied markers (except for *BRCA1*) on the panel of breast cancer cell lines.

| Breast cancer cell lines | EMP-unassigned group |  |  |  |  |  |  | Mesenchymally-expressed group |  |  |  |  | Epithelially-expressed group |  |  |
| --- | --- | --- | --- | --- | --- | --- | --- | --- | --- | --- | --- | --- | --- | --- | --- |
|  | *AKR1B1* | *APC* | *RASSF1A* | *RARβ* | *SFRP2* | *GFRA1* | *CDKN2A* | *Twist1* | *CRABP1* | *DKK3* | *EGFR* | *VIM* | *GRHL2* | *MIR200C* | *CDH1* |
| MFM-223 | very high | 100% | 100% | 100% | 10-50% | 100% | moderate | high | 100% | 100% | low | very low | 0 | 0 | 0 |
| MDA-MB-453 | very high | 100% | moderate | 100% | 50-100% | high | 0 | high | moderate | 100% | 10-50% | 100% | 0 | 0 | 0 |
| BT-474 | 100% | 10-50% | 100% | 100% | 50-100% | 0 | 0 | very low | high | moderate | very low | 0 | 0 | 0 | 0 |
| ZR-75-1 | moderate | 100% | 100% | high | moderate | very low | low | moderate | low | low | very low | moderate | 0 | 0 | 0 |
| HCC1419 | very high | 0 | 100% | very low | high | 0 | 0 | very low | high | moderate | very low | moderate | 0 | 0 | 0 |
| UACC-893 | very high | 50% | 100% | high | high | moderate | 0 | high | low | moderate | 0 | low | 0 | 0 | 0 |
| SK-BR-3 | moderate | low | 50% | high | moderate | 0 | 0 | very low | 0 | 0 | very low | moderate | 0 | 0 | n.a. |
| T-47D | moderate | 0 | 100% | very low | moderate | 0 | 100% | high | moderate | moderate | moderate | 100% | 0 | 0 | 0 |
| CAL-148 | very high | 100% | 100% | high | moderate | 0 | 0 | 100% | high | low | very low | moderate | 0 | 0 | 0 |
| MCF-7 | very high | 10-50% | 100% | 100% | high | 0 | n.a. | high | moderate | moderate | low | 100% | 0 | 0 | 0 |
| UACC-732 | high | 50% | 100% | high | moderate | 0 | 0 | high | low | moderate | very high | 100% | 0 | 0 | 0 |
| MDA-MB-468 | 0 | 3% | 100% | 50-100% | moderate | moderate | 0 | high | low | moderate | 0 | 0 | 0 | 0 | 0 |
| HCC1954 | 100% | 100% | 100% | high | high | low | 100% | moderate | moderate | high | very low | 0 | 0 | 0 | 0 |
| HCC70 | 0 | 0 | 0 | 50% | very low | low | 0 | moderate | 0 | moderate | very low | 0 | 0 | 0 | 0 |
| BT-20 | 0 | 100% | 50% | 100% | 100% | 100% | n.a. | 100% | high | 100% | 0 | 100% | 0 | 0 | 0 |
| HCC1806 | 0 | 0 | 0 | very low | 0 | 0 | n.a. | 0 | very low | 0 | very low | very low | 0 | 0 | 0 |
| HCC1569 | 10-50% | 50-100% | 10-50% | very low | 100% | 100% | low | 100% | 0 | moderate | very low | 0 | 0 | 0 | 0 |
| SUM-149PT | 0 | 0 | 0 | moderate | low | 0 | n.a. | very low | very low | very low | very low | 0 | 0 | 0 | 0 |
| CAL-120 | 0 | 0 | 0 | 50% | moderate | 0 | 0 | very low | low | 0 | very low | 0 | very low | very low | very low |
| MCF 10A | 0 | 0 | moderate | high | low | low | n.a. | very low | very low | low | 0 | 0 | high | 0 | 0 |
| PMC42-LA | 0 | 0 | moderate | 100% | low | 0 | n.a. | low | very low | 0 | 0 | 0 | 10% | low | 0 |
| PMC42-ET | 0 | 0 | moderate | 50-100% | moderate | 0 | n.a. | moderate | low | 0 | 0 | 0 | 50-100% | high | very low |
| MDA-MB-231 | 0 | 0 | 50% | very low | high | moderate | n.a. | 100% | low | high | very low | 10% | moderate | moderate | moderate |
| BT-549 | 0 | 0 | very low | very low | 0 | 0 | 0 | low | low | very low | very low | 0 | high | moderate | very low |
| SUM-159PT | 0 | 0 | 50% | moderate | high | low | very low | high | 0 | 0 | very low | 0 | high | high | moderate |
| Hs578T | 0 | 0 | 10-50% | 0 | very low | n.a. | n.a. | high | 0 | 0 | 0 | 0 | moderate | high | moderate |

NB: Cell lines were ranked in the epithelial-mesenchymal spectrum from the epithelial luminal (top), intermediate (middle) to mesenchymal basal (bottom) groups according to Tan *et al*. [15]. The levels of homogeneous methylation were stated numerically as percentage of methylation. The estimated levels of heterogeneous methylation were described as ‘very high’, ‘high’, ‘moderate’, ‘low’ and ‘very low’. Zero means no methylation. ‘n.a.’ means no amplification.

### A generally defined yet complex pattern of methylation in breast cancer cell lines

Since methylation at the promoter region generally silences gene expression, methylation of markers that are typically expressed in epithelial cells, i.e., *GRHL2, MIR200C*, and *CDH1*, were considered potentially mesenchymal-specific, and methylation of markers that are typically expressed in mesenchymal cells, i.e., *TWIST1, DKK3, VIM, CRABP1* and *EGFR*, were considered potentially epithelial-specific. In addition, we assessed several DNA methylation markers that have not yet been associated with epithelial or mesenchymal status, but have been proposed for monitoring of MRD (*AKR1B1, APC, RASSF1A, RARß, SFRP2* and *GFRA1)*, as well as two breast cancer tumour suppressor gene markers (*BRCA1* and *CDKN2A*).

In Table 3, based on gene expression profiles, the genes were grouped into epithelially-expressed (epithelial group), mesenchymally-expressed (mesenchymal group) and those for which epithelial or mesenchymal status remains to be determined (EMP-unassigned group). The methylation result of each cell line for each gene marker is colour-coded with red being 100% homogeneous methylation or very high heterogeneous methylation, orange being 50% homogeneous methylation or high heterogeneous methylation, yellow being 10-50% homogeneous methylation or moderate heterogeneous methylation, light yellow being 10% homogeneous methylation or low heterogeneous methylation and white being less than or equal to 3% homogeneous methylation or no methylation or very low heterogeneous methylation. To compensate for non-specific background, methylation levels less than 3% for homogeneous methylation and very low heterogeneous methylation were ignored, although they may still represent true minor cellular sub-populations consistent with cellular plasticity.

The mesenchymally-expressed markers *DKK3, CRABP1, VIM* and *EGFR* are methylated at moderate to high levels in the majority of epithelial cells. Several epithelial cell lines were fully methylated for these genes, for example MFM-223 for *CRABP1*, MFM-223 and MDA-MB-453 for *DKK3*. The methylation of *DKK3* and *CRABP1* is not restricted to epithelial cell lines but is also seen in some intermediate and mesenchymal basal cell lines (Table 3)*. DKK3* was methylated in the epithelial cell lines BT-20, MCF-7, T-47D and ZR-75-1, in the mesenchymal cell line MDA-MB-231 but not in Hs578T, similar to the findings of Veeck *et al*. [27]. SK-BR-3 is the only epithelial cell line which is not methylated for *DKK3*.

In contrast, *VIM* is methylated in most epithelial cell lines in our study except for BT-474. While the majority of intermediate and mesenchymal basal cell lines are unmethylated for *VIM*, the BT-20 and MDA-MB-231 cell lines are methylated at 100% and 10%, respectively. Interestingly, the archetypal mesenchymal cell line MDA-MB-231 displays moderate to high methylation levels for the mesenchymal markers *DKK3, CRABP1* as well as *VIM*, in contrast to other mesenchymal cell lines, which may indicate a higher propensity for plasticity of this cell line.

*EGFR* is methylated in only few epithelial cell lines compared to other mesenchymal markers. Three epithelial cell lines, i.e. MDA-MB-453, T-47D and UACC-732, show relatively moderate to high methylation level with UACC-732 being methylated at a high level for *EGFR*. The methylation of *EGFR* in MDA-MB-453 is consistent with the study of Montero et al. [28]. All the mesenchymal cell lines have no or very low methylation for this marker, consistent with its expression in mesenchymal cell lines.

*TWIST1* is methylated across the epithelial and mesenchymal spectrum from low to very high level. The methylation distribution of *TWIST1* is different from other mesenchymally-expressed markers, the methylation of which is predominantly in epithelial cell lines.

*RARß* is also methylated in both epithelial and mesenchymal cell lines, with the methylation level being higher in epithelial cell lines except for HCC1419 and T-47D. Most mesenchymal cell lines are also methylated at moderate to high levels for *RARß* with the exceptions of Hs578T having no methylation, and MDA-MB-231 and BT-549 having very low level of methylation.

Similar to the methylation distribution of *RASSF1A* and *RARß, SFRP2* was methylated across the epithelial and mesenchymal spectrum, except for a few cell lines including HCC70, HCC1806, BT-549 and Hs578T. Its methylation level was moderate to high in the majority of cell lines, in both epithelial and mesenchymal states. Surprisingly, the intermediate cell lines BT-20 and HCC1569 displayed the highest methylation level of 100% for *SFRP2*.

*GFRA1* was methylated in fewer epithelial, intermediate and mesenchymal cell lines, compared to other genes in the EMP-unassigned group. Interestingly, it was also fully methylated in the intermediate cell lines BT-20 and HCC1569 as seen for *SFRP2* methylation, and the epithelial cell line MFM-223.

The other cancer-associated markers *APC* and *AKR1B1* show relatively high methylation level in only epithelial cell lines whilst no methylation is observed in mesenchymal cell lines. A few exceptions for *APC* are the epithelial cell lines SK-BR-3 with low methylation level, and HCC1419 and T-47D with no methylation for this marker.

The majority of cell lines did not amplify for *CDKN2A*. Among those, MCF-7, MDA-MB-231, BT-20 and Hs578T were reported with homozygous deletion by Paz *et al*. [29]. T-47D and HCC1954 cell lines are homogeneously methylated at 100%, MFM-223 cell line is moderately methylated and ZR-75-1 and HCC1569 are methylated at low levels for *CDKN2A*. All the breast cancer cell lines were unmethylated for *BRCA1*.

Unsupervised hierarchical clustering of all the breast cancer cell lines for all the gene markers gave rise to four main clusters (Figure 3). The methylation profiles separated the cell lines according to epithelial and mesenchymal phenotypes, and also the molecular subtypes. Cluster 1 contained predominantly epithelial cell lines of luminal subtype. Cluster 2 was comprised of both luminal and basal A cell lines. Cluster 3 had mainly basal A intermediate cell lines while cluster 4 was comprised of all the mesenchymal basal B and triple negative breast cancer cell lines. Most ER+, PR+ and HER2+ cell lines fell into clusters 1 and 2.

**Figure 3:**
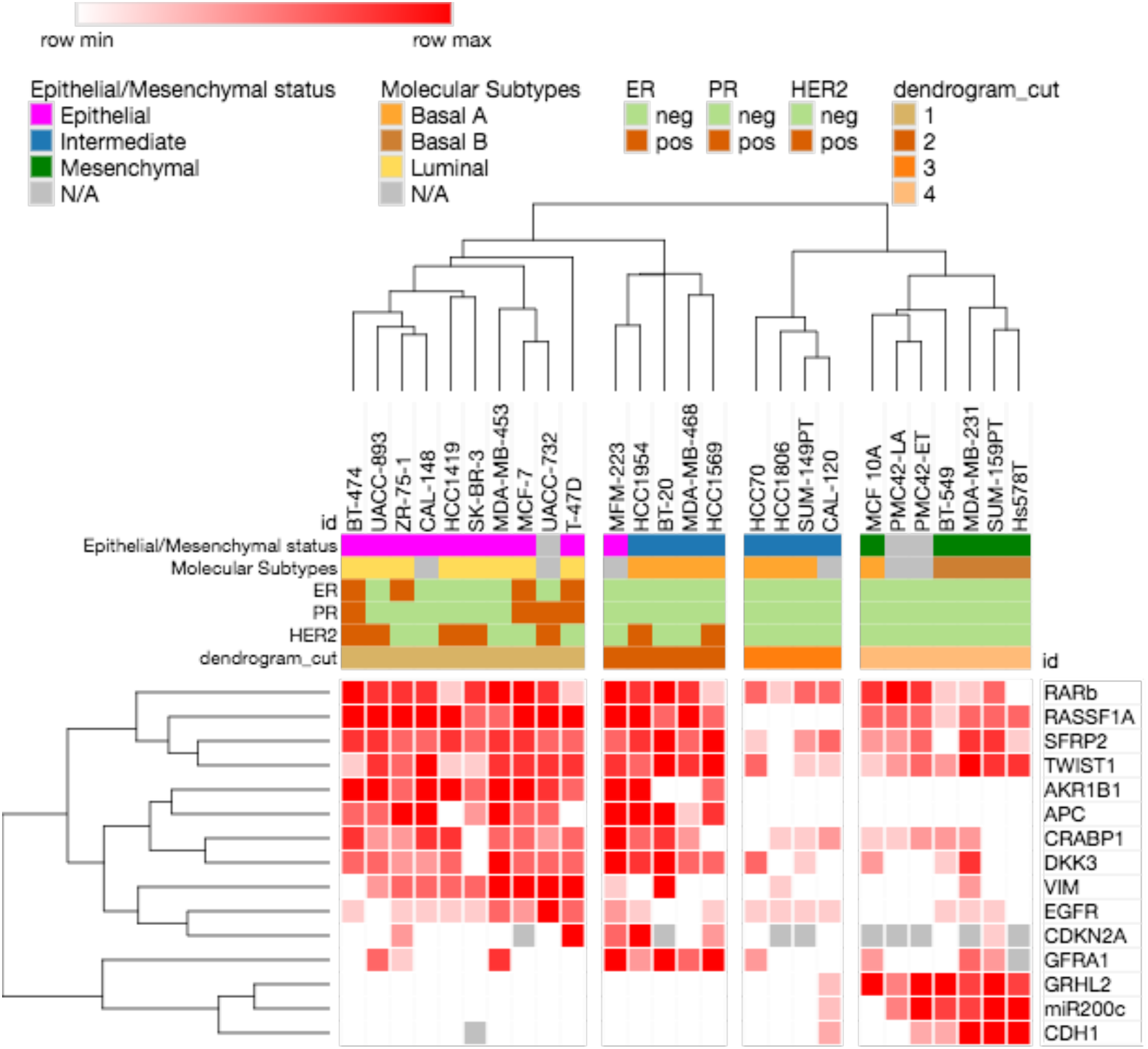
Unsupervised hierarchical clustering of the breast cancer cell lines according to methylation the selected gene markers.

## Discussion

In this study, we determined the methylation status of sixteen gene markers in a panel of twenty-six breast cancer cell lines spanning the epithelial-mesenchymal spectrum. This knowledge may be of particular relevance for selecting an optimal panel of methylation markers to assess MRD in breast cancer, given that ctDNA may derive from tumour cells across the epithelial-mesenchymal spectrum. Our study also provides some insight on the role of DNA methylation in the molecular stratification of tumours.

A panel of methylation-based markers, whose expression is associated with mesenchymal or epithelial phenotypes, were selected and grouped into mesenchymally-expressed (i.e., *TWIST1, DKK3, VIM, CRABP1* and *EGFR*) and epithelially-expressed (i.e., *GRHL2, MIR200C*, and *CDH1*) markers, respectively. Methylation of most promoter regions is associated with lack of gene expression. Thus, when a gene is predominantly expressed in mesenchymal cells, its methylation would be expected to be predominantly epithelial. Our results showed a close-to-expected distribution of the epithelial methylation and mesenchymal methylation markers along the epithelial-mesenchymal spectrum.

The majority of mesenchymal methylation markers were predominantly methylated in epithelial cell lines, while the epithelial methylation markers were methylated in mesenchymal cell lines. The status of mesenchymal methylation markers is thus distinctly different between epithelial and mesenchymal cell lines. All the epithelial and intermediate cell lines are unmethylated for *GHRL2, MIR200C* and *CDH1*, whereas all the mesenchymal cell lines are methylated at moderate to high levels for *GHRL2* and *MIR200C*.

Three mesenchymal cell lines MDA-MB-231, SUM-159PT and Hs578T are methylated at moderate levels for *CDH1*, consistent with Lombaerts *et al.*, who also showed partial methylation in mesenchymal breast cancer cell lines [30]. SK-BR-3 did not show amplification in our assay for *CDH1*, consistent with the report of a promoter deletion by Lombaerts *et al*. [30].

*GRHL2* is one of the most commonly studied EMT markers, and its expression is usually elevated in epithelial luminal cells and down-regulated in mesenchymal breast cancer cell lines [31–33]. Here, we showed that epithelial breast cancer cells have no methylation whereas mesenchymal basal cell lines displayed moderate to high methylation levels for this marker. This is consistent with findings of the differential methylation between epithelial and mesenchymal cell lines in non–small cell lung cancer [34]. The negative correlation between gene expression and methylation for *GRHL2* in our breast cancer cell line panel suggests gene expression is regulated by methylation.

We showed that *MIR200C* was methylated at moderate to high level in mesenchymal breast cancer cell lines. Similarly, Davalos *et al*. showed methylation of the CpG island associated with the MiR-200 cluster including *MIR200C* in mesenchymal MDA-MB-231 cells, whereas this was unmethylated in epithelial MCF-7 cells [35]. Neves *et al*. demonstrated that the expression of the miR200c/141 cluster was regulated by DNA methylation, and that the hypermethylation of the *MIR200C* promoter region was tightly correlated with mesenchymal phenotype and invasive capacity in a panel of eight breast cancer cell lines [36]. In addition, the loss of MiR-200 expression was found in the stem cell-like fractions from metastatic breast cancer, and miR200c/141 cluster repression by DNA methylation was associated with the transition from non-stem to a stem-like phenotype in human mammary epithelial (HMLE) cells [37].

Another interesting observation from this study is the very restricted methylation status of *EGFR*. *EGFR* expression is often higher in mesenchymal basal cell lines [2, 24], and thus was expected to be predominantly methylated in epithelial luminal cell lines. However, unlike other epithelially methylated markers, *EGFR* is only methylated at moderate to high levels in two HER2-amplified epithelial cell lines, i.e., MDA-MB-453 and UACC-732, and the HER2-normal epithelial cell line T-47D (Table 1). Interestingly, the HER2-amplified cell lines MDA-MB-453 and UACC-732 were resistant to the pan-HER receptor tyrosine kinase inhibitor dacomitinib, which selectively targets EGFR, HER2, and HER4, whereas the majority of HER2-amplified cell lines were sensitive to the drug [25]. The HER2-normal T-47D cell line [23, 38] also showed resistance to dacomitinib [25]. It can be speculated that *EGFR* methylation and its consequent absence of expression could be one of the resistance mechanisms to EGFR inhibitors in breast cancer.

DNA methylation markers previously used in monitoring MRD, including the commonly used *RASSF1A, APC, TWIST1* and *RARß*, and the less commonly used *AKR1B1, SFRP2, GFRA1, BRCA1* and *CDKN2A*, were also assessed, as the knowledge of how their methylation relates to epithelial-mesenchymal status is relevant to their use as MRD markers.

*RASSF1A* methylation was distributed widely across the epithelial-mesenchymal spectrum. Promoter methylation of the *RASSF1A* gene is the most common cause of the gene’s inactivation, and is associated with tumour invasion and metastasis [39, 40]. This is concordant with the observed methylation of the gene in many cell lines across the epithelial-mesenchymal spectrum in our study. We also observed that the methylation level is higher in epithelial cell lines and most epithelial luminal cell lines have complete methylation. This might be related to the report that *RASSF1A* hypermethylation was significantly more common in ER-positive and HER2-positive tumours [41].

*TWIST1* promoter methylation has been reported to be a useful marker to detect breast cancer [42]. *TWIST1* showed methylation across the epithelial-mesenchymal spectrum. From an MRD perspective, a marker that is methylated across the EMP spectrum will be useful. Figure 3 shows that it clusters with *RASSF1A, RARß* and *SFRP2*. Although *TWIST1* is considered a master regulator of EMT, DNA methylation may not be the main mechanism that regulates *TWIST1* expression in mesenchymal cells. Indeed, it has been shown that *TWIST1* methylation did not correlate with either mRNA or protein expression in tumours [42].

*RARß* was methylated at some level in every cell line except Hs578T. Although its methylation was very low in a subset of both epithelial and mesenchymal cell lines, it still proved to be one of the most useful markers in terms of coverage across the EMT spectrum.

*SFRP2* also showed a distribution of methylation across the epithelial-mesenchymal spectrum. [43]. Its methylation has been detected at high frequency in cancer patients, typically in lung cancer and gastric cancer [44–46]. It is a canonical Wnt pathway inhibitor, and has been reported to be a promising novel therapeutic marker for several tumor types [43, 45, 47]. The loss of *SFRP2* expression interferes with the growth of mammary epithelial cells [43]. *SFRP2* was found to promote epithelial cell transformation and to stimulate cell adhesion to the extracellular matrix in breast tumours, which consequently promotes resistance to apoptosis [48].

*APC* and *AKR1B1* are methylated only in the epithelial cell lines and in some intermediate cell lines. *APC* is an inhibitor of the Wnt signaling pathway that acts by inactivating ß-catenin [49]. Its loss of expression and the up-regulation of ß-catenin were first described in the development of colorectal cancer and then in many other cancers [50, 51]. Wnt signaling has been implicated in EMP [52], consistent with the selective methylation of *APC* in epithelial cell lines. *AKR1B1* has also been shown to be involved in EMT where its over-expression is associated with up-regulation of EMT markers in lens epithelial cells [53], and the selective absence of *AKR1B1* methylation in mesenchymal cell lines consistent with the potential role in breast cancer EMT recently reported in basal breast cancers [54].

Overall, the methylation data confirmed the general distinction between epithelial and mesenchymal cell lines, with some exceptions showing the complex distribution of methylation in the epithelial-mesenchymal spectrum. By looking at the methylation status of the markers in Table 3, it is possible to distinguish between epithelial cell lines and mesenchymal cell lines. Epithelial cell lines are methylated for the majority of previously used cancer markers in MRD and mesenchymal markers, whereas mesenchymal cell lines are methylated for the epithelially-expressed markers.

Epithelial cell lines display higher methylation levels than mesenchymal cell lines, and the methylation seen in the mesenchymal cell lines is often heterogeneous. This observation in cell lines reflects the results from recent studies showing that luminal breast tumours have overall higher methylation level than other subtypes [55–57]. The lower methylation levels of these epithelial markers is perhaps explained by the previous observation that mesenchymal basal cell lines seem to repress genes largely by histone H3K27 methylation in preference to DNA methylation [58, 59], and that H3K27 methylation is associated with lower level of DNA methylation [60].

Some interesting patterns were seen for the markers that have been commonly used to monitor MRD. Almost all epithelial cell lines are fully methylated for *RASSF1A* at high levels whilst most mesenchymal cell lines are also methylated, but to a lesser extent. However, cell lines with an intermediate epithelial-mesenchymal phenotype show a more complex methylation status, and the mesenchymal cell line BT-549 and most of the intermediate cell lines, including HCC70, HCC1806, SUM-149PT and CAL-120, being unmethylated for *RASSF1A*.

Cell lines with intermediate epithelial-mesenchymal phenotypes show a more complex methylation status of the studied genes, which might reflect their plasticity. A recent study by Chung *et al*. [59]. suggested that cells with the intermediate epithelial phenotype have a less tight control of genes and as a consequence more plasticity than the epithelial cells in ovarian cell type.

By taking advantage of MS-HRM to visualize the methylation patterns based on the melting profiles, our study revealed that only the markers used commonly to monitor MRD (i.e., *APC, RASSF1A* and *RARß*) displayed homogeneous methylation overall, while the majority of the other markers showed heterogeneous methylation (Table 3). Previous genome-wide methylation studies have shown that the DNA methylation profiles of breast cancer cell lines largely mirror those of primary tumours [61, 62]. Based on the methylation in breast cancer cell lines, it can be predicted that tumour samples may also predominantly show heterogeneous methylation for these studied genes.

Genes with homogeneous methylation are more easily detected compared to those with heterogeneous methylation, especially in the MRD context, due to the simultaneous methylation at each CpG. Our observation in terms of homogeneous and heterogeneous methylation profiles might help to explain why the methylation of *APC, RASSF1A* and *RARß* genes with homogeneous pattern have been detected at high frequency, and used successfully in detecting various tumour types, and also MRD. Therefore, we consider that homogeneously methylated markers are more useful for the detection and monitoring of MRD. Our study also indicates that the visualization and identification of the methylation profiles are essential for marker selection in future methylation studies.

It remains unclear whether specific mesenchymally methylated markers need to be developed. The majority of cells in most breast tumours are epithelial and thus should contribute to the majority of ctDNA. In addition, the majority of epithelially-expressed (mesenchymally methylated) markers are methylated at varying frequencies in peripheral blood DNA, which makes the major contribution to cell free DNA and confounds the methylation-based detection of ctDNA. Thus, markers such as *RASSF1A, RARß, TWIST1* and *SFRP2*, whose methylation covers the epithelial-mesenchymal spectrum are likely to be the most useful for monitoring ctDNA.

## Conclusions

This is the first study to describe locus-specific DNA methylation status associated with the epithelial-mesenchymal spectrum. We determined the methylation status of a panel of genes in breast cancer cell lines across the epithelial-mesenchymal spectrum. This included those genes that might be expected to be epithelial or mesenchymal markers, in addition to established DNA methylation markers for monitoring MRD. We have shown that the methylation of many of these markers was associated with the epithelial or mesenchymal status of the cell lines whereas others are less specific. In addition, the methylation patterns generally support the previously known gene expression–based classification of epithelial/mesenchymal status. Cancer markers previously used to detect MRD are predominantly methylated in epithelial and intermediate cell lines with *RASSF1A, TWIST1* and *RARß* methylation covering more of the epithelial-mesenchymal spectrum in the cell line panel. A similar approach can be used to assess further markers in order to develop better DNA methylation markers for the detection of ctDNA in breast cancer.

## Additional files

**Additional file 1: Supplementary Figure 1. MS-HRM melt profiles and bisulfite pyrosequencing data of representative samples for (A, B) *RARß*, (C, D) *CRABP1*, (E, F) *AKR1B1*, (G, H) *SFRP2* and (I, J) *GFRA1***. A standard methylation series was included in both MS-HRM and pyrosequencing experiments as a guide for determining methylation and for calling methylation levels. (A, C, E, G, I) MS-HRM melt profiles show homogeneous methylation and heterogeneous methylation. Methylation level was numerical for samples with homogeneous methylation, whereas it was called ‘very high’, ‘high’, ‘moderate’, ‘low’ and ‘very low’ for very high level, high level, moderate level, low level and very low level of heterogeneous methylation, respectively. (B, D, F, H, J) Pyrosequencing data validated the results from MS-HRM and provided complementary information on methylation level at each CpG position.

**Additional file 2: Table S1.** Oligonucleotide sequences for methylation-sensitive high-resolution melting (MS-HRM) assays and characteristics of the corresponding amplicons. (CpG sites in bold, converted Cs as capital Ts or As).

**Additional file 3: Table S2.** Summary of PCR conditions for sixteen methylation-sensitive high-resolution melting (MS-HRM) assays.

**Additional file 4: Table S3.** Sequences before bisulfite treatment, sequences to analyse and dispensation order for tested bisulfite pyrosequencing assays. (bold nucleotides in each dispensation order are control nucleotides added by the Qseq advanced software (Bio Molecular Systems, v2.0.11), which allow the detection of incomplete bisulfite conversion or non-specific products).

## Abbreviations

EMP: epithelial-mesenchymal plasticity; EMT: epithelial-mesenchymal transition; MET: mesenchymal-epithelial transition; MS-HRM: methylation-sensitive high resolution melting; PCR: polymerase chain reaction; MSP: methylation-specific PCR; CpG: cytosine-guanine dinucleotides; MRD: minimal residual disease; CTCs: circulating tumour cells; ctDNA: circulating tumour DNA; ER: estrogen receptor; PR: progesterone receptor; HER2: human epidermal growth factor receptor 2; ATCC: American Type Culture Collection; AKR1B1: Aldo-Keto Reductase Family 1, Member B1; APC: Adenomatous Polyposis Coli; RASSF1A: Ras Association Domain Family Member 1A; RAR*ß*: Retinoic Acid Receptor Beta; CDKN2A: Cyclin-Dependent Kinase Inhibitor 2A; BRCA1: Breast Cancer 1; TWIST1: Twist Family BHLH Transcription Factor 1; CRABP1: Cellular Retinoic Acid Binding Protein 1; DKK3: Dickkopf WNT Signaling Pathway Inhibitor 3; EGFR: Epidermal Growth Factor Receptor; VIM: Vimentin; GRHL2: Grainyhead Like Transcription Factor 2; MIR200C: MicroRNA 200c; CDH1: Cadherin 1; SFRP2: Secreted Frizzled Related Protein 2; GFRA1: GDNF Family Receptor Alpha 1.

## Authors’ contribution

AL performed the methylation assays, analysed, interpreted the data and prepared the manuscript. MS performed methylation assays of *SFRP2* and *GFRA1*, provided additional figures. TZT and JPT provided the EMT scores for the studied cell lines and Figure 1. EWT helped to interpret and put the results into context, and critically reviewed the manuscript. AD conceived the study and provided overall study guidance, interpreted the data, put the results into context and critically reviewed the manuscript. All the authors read and approved the manuscript.

## Acknowledgements

This work was supported by the above funding bodies. The authors would also like to acknowledge Dr. Mark Waltham for providing DNA from PMC42-ET and PMC42-LA cell lines, Riley Morrow and Tracy Cardwell for providing cell line pellets whose DNA was extracted and used in this study.

